# A long-chain fatty acid elongase Elovl6 regulates mechanical damage–induced keratinocyte death and skin inflammation

**DOI:** 10.1101/264838

**Authors:** Yoshiyuki Nakamura, Takashi Matsuzaka, Satoko Tahara-Hanaoka, Kazuko Shibuya, Hitoshi Shimano, Chigusa Nakahashi-Oda, Akira Shibuya

## Abstract

Mechanical damage on the skin not only affect the barrier function but also induce various immune responses, which trigger or exacerbate the inflammation in healthy individuals and patients with inflammatory skin diseases. However, how mechanical damage-induced skin inflammation is regulated remains largely unknown. Here, we show that mechanical damage due to tape stripping triggered keratinocyte death and release of danger-associated molecular patterns (DAMPs) such as high-mobility group box 1 protein (HMGB-1) and IL-1α, which induced production of proinflammatory cytokines and chemokines IL-1β and CXCL-1 by keratinocytes in mice. We also show that a long-chain fatty acid elongase Elovl6 is expressed in keratinocytes. Mice deficient in Elovl6 had increased epidermal levels of cis-vaccenic acid (CVA); this accelerated keratinocyte death triggered by tape stripping and release of DAMPs and exacerbated skin inflammation. Our results demonstrate that Elovl6 regulates mechanical damage–triggered keratinocyte death and skin inflammation.

## Introduction

The mechanical damage induced by physical forces—including stretching, compression, and friction—on epithelial and endothelial cells plays a critical role in tissue homeostasis (Abe and Berk 2014; Angelini *et al*. 2012; Hofmann *et al*. 2004; Reichelt 2007; Wyatt *et al*. 2016). Under physiologic conditions, keratinocytes are the epidermal cell population most affected by mechanical damage (Reichelt 2007), which induces them to proliferate and produce cytokines. Streching of keratinocytes in vitro opens Ca^2+^ channels, resulting in the phosphorylation of Akt (Yano *et al*. 2006). This process also induces clustering and co-localization of β-integrins and epidermal growth factor receptor, followed by activation of the downstream signaling molecule extracellular signal-regulated kinase (ERK) 1/2 (Knies *et al*. 2006).

Mechanical damage not only affect the barrier function of the skin but also induce various immune responses (Verhoeven *et al*. 2008), which trigger inflammation at the site of the stress on the skin of healthy individuals. Moreover, mechanical damage on the skin exacerbates the inflammation in patients with inflammatory skin diseases. For example, scratching of itching lesions exacerbates the skin inflammation in atopic dermtitis (AD), which is called the itch-scratch cycle (Verhoeven *et al*. 2008; Verhoeven *et al*. 2009; Wahlgren 1999). In addition, scratching induces development of new skin lesions in psoriasis, well known as the Koebner phenomenon (Köbner 1876). However, how mechanical damage-induced skin inflammation is regulated remains largely unknown.

Elongation of long-chain fatty acids family member 6 (Elovl6) is a rate-limiting microsomal enzyme that catalyzes the elongation of saturated and monounsaturated fatty acids (Saito *et al*. 2011). Elovl6 elongates palmitate (PA) (C16:0) to stearate (SA) (C18:0) and palmitoleate (POA) (C16:1n-7) to cis-vaccenic acid (CVA) (C18:1n-7) (Saito *et al*. 2011). Elovl6 is highly expressed in white adipose tissue and liver (Matsuzaka *et al*. 2002). In previous studies, mice deficient in Elovl6 (*Elovl6^-/-^* mice) had increased levels of PA in the liver and lung (Matsuzaka *et al*. 2007; Sunaga *et al*. 2013). Elovl6 is involved in metabolic diseases, such as insulin resistance (Matsuzaka *et al*. 2007) and atherogenesis (Saito *et al*. 2011), as well as inflammatory diseases, including attenuated high-fat–diet-induced hepatic inflammation (Matsuzaka *et al*. 2012) and regulated bleomycin-induced pulmonary fibrosis (Sunaga *et al*. 2013). In addition, Elovl6 is highly expressed in skin (Matsuzaka *et al*. 2002), which is one of the most lipid-enriched organs. Lipids in the skin play crucial roles in homeostasis; they are involved in epidermal permeability and barrier function (Ishikawa *et al*. 2010), the composition of microbiota (Nguyen *et al*. 2016), epithelialization (Liu *et al*. 2014), and inflammation (Zhang *et al*. 2015).

In the current study, we examined how mechanical damage induces skin inflammation and whether long-chain fatty-acid composition regulated by Elovl6 is involved in mechanical damage onto the skin.

## Results

### *Elovl6^-/-^* mice show exacerbated mechanical damage–induced skin inflammation

Tape stripping, which mimics scratching, is a well-established method for inducing mechanical stress or damage on the skin (Onoue *et al*. 2009; Takahashi *et al*. 2013). To investigate the role of Elovl6 in mechanical damage–induced skin inflammation, we established a mouse model of dermatitis by using repeated tape stripping twenty times, which induced the skin damage and barrier disruption. After this treatment, erythema was more severe in *Elovl6^-/-^* mice than in wild-type mice (Figure 1A). Moreover, the epidermis was thicker, and neutrophil infiltration was significantly greater, in *Elovl6^-/-^* mice than in wild-type mice (Figure 1B–D). Since *Elovl6* expression was higher in the epidermis than in the dermis (Figure supplement 1A), we speculated that Elovl6 is expressed in keratinocytes. Indeed, the epidermis in mice deficient in Elovl6 specifically in the keratinocytes (*Elovl6*^fl/fl^ *K14-Cre* mice) showed significantly decreased Elovl6 expression (Figure supplement 1B). As in *Elovl6^-/-^* mice, *Elovl6*^fl/fl^ *K14-Cre* mice also showed increased epidermal thickness and neutrophil infiltration after tape stripping compared with control mice (Figure 1E, F).

**Fig. 1.**
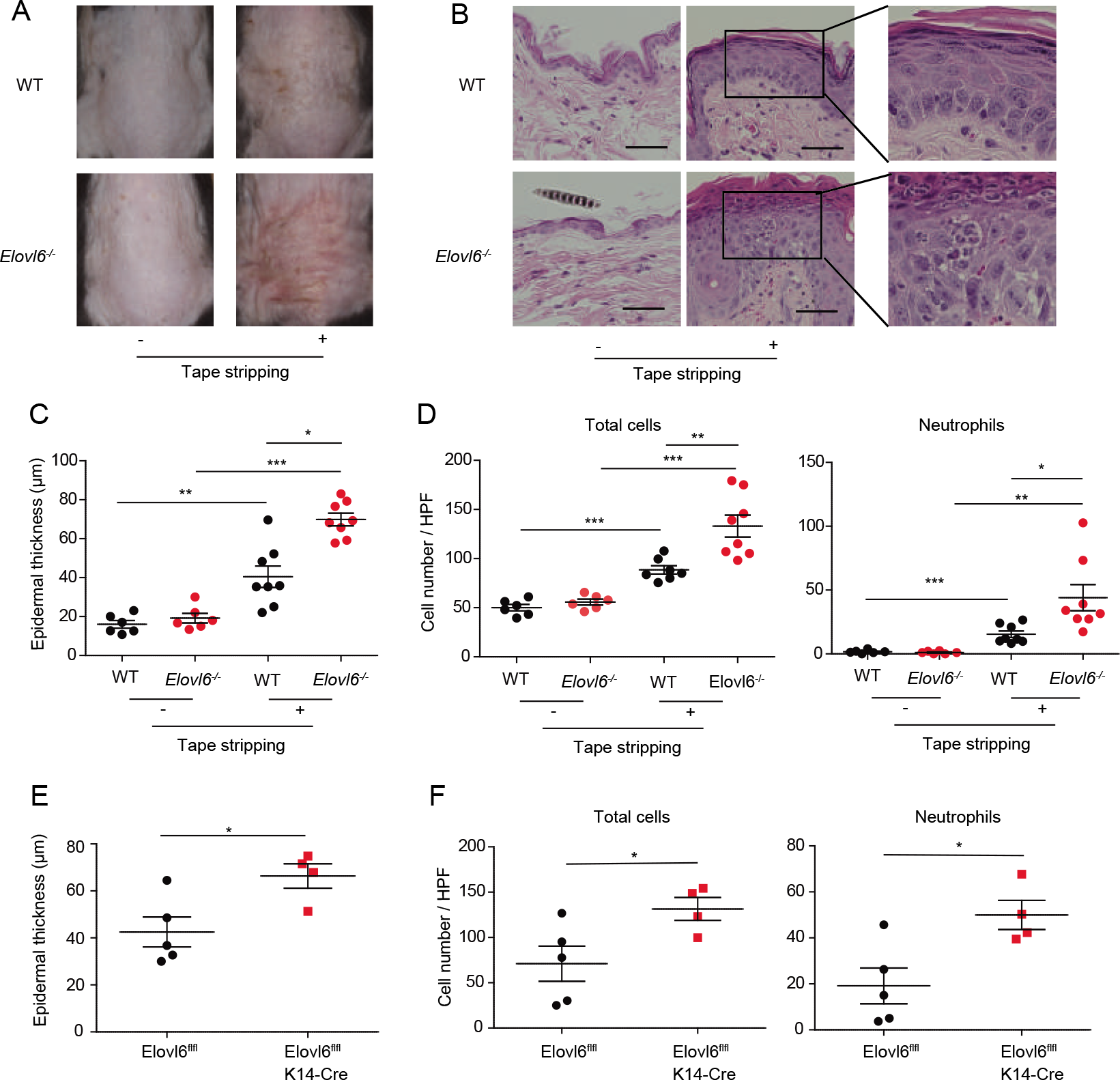
*Elovl6^-/-^* mice show exacerbation of dermatitis. (A to D) Representative gross findings (A), histology (hematoxylin and eosin staining) (B), epidermal thickness (C), and numbers of total infiltrating cells and neutrophils in the dorsal skin (D) of wild-type (n =6 or 8) and *Elovl6^-/-^* (n =6 or 8) mice before and on day 9 after the start of tape stripping. (E, F) Epidermal thickness (E) and numbers of total infiltrating cells and neutrophils in the skin (F) of *Elovl6^fl/fl^* (n = 5) and *Elovl6*^fl/fl^ *K14*-Cre (n = 4) on day 9 after the start of tape stripping. Black bars indicate scale (50 μm) (B). Error bars indicate 1 SD; *, *P*< 0.05; **, *P* < 0.01; ***, *P* < 0.001. Data are representative of three (A-D) and two (E, F) independent experiments.

To address how Elovl6 suppressed mechanical damage-induced skin inflammation, we compared skin barrier function of *Elovl6^-/-^* mice with that of wild-type mice. However, it was comparable between two genotypes of mice, as determined by toluidine blue skin permeability assay and a transepidermal water loss test (Sassa *et al.* 2013) (Figure supplement 2A, B). These results suggested that Elovl6 in the keratinocytes suppressed mechanical damage–induced skin inflammation, in which a novel mechanism might be involved.

### *Elovl6^-/-^* mice show increased IL-1β and CXCL-1 production after mechanical damage

To investigate how Elovl6 suppressed mechanical damage-induced skin inflammation, we examined the expression levels of pro-inflammatory and anti-inflammatory cytokines and chemokines potentially involved in dermatitis (Effendy *et al*. 2000). Among them, transcript levels of *Il1b* and *Cxcl1* in epidermis were increased in both wild-type and *Elovl6^-/-^* mice after tape stripping (Figure 2A and Figure supplement 3). Moreover, *Elovl6^-/-^* mice showed higher expression of *Il1b* and *Cxcl1* than did wild-type mice after tape stripping (Figure 2A). In accordance with these results, the concentrations of IL-1β and CXCL-1 were significantly higher in the culture supernatants of *Elovl6^-/-^* epidermis harvested from mice after tape stripping than in those from wild-type epidermis (Figure 2B). These cytokine levels in the epidermis from mice deficient in an adaptor of Toll-like receptors (TLRs) MyD88, but not TRIF, were lower than those in wild-type mice after tape stripping (Figure 2C). These results suggest that Elovl6 suppressed mechanical damage-induced IL-1β and CXCL-1 productions that are dependent on MyD88.

**Fig. 2.**
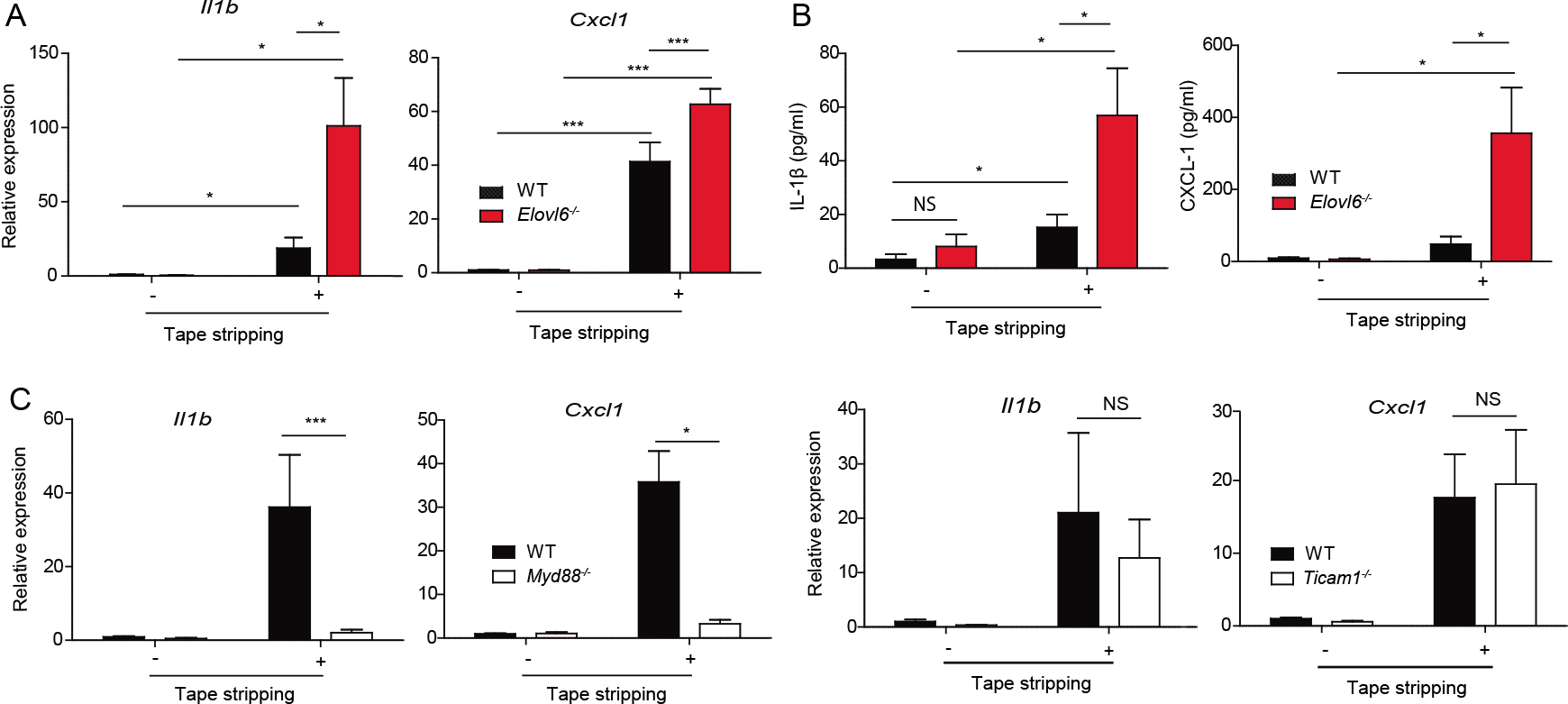
Elovl6 suppresses mechanical damage–induced IL-1β and CXCL-1 production. (A) Quantitative RT-PCR analysis of epidermis of wild-type and *Elovl6^-/-^* mice isolated 6 h after tape stripping (n = 10 in each group). (B) Epidermis was isolated before, and 12 h after, tape stripping from wild-type and *Elovl6^-/-^* mice and cultured for 24 h. The concentrations of IL-1β and CXCL-1 in the supernatants were measured by using cytometric bead array (n = 11 per group). (C) Quantitative RT-PCR analysis of *¡lift* and *Cxcl1* in the epidermis of wild-type, *Myd88^-/-^*, and *Ticami^-/-^* mice before, and 6 h after, tape stripping (n = 7 to 11 per group). Error bars indicate 1 SD; *, *P*< 0.05; ***, *P* < 0.001. Data are representative of three independent experiments.

### *Elovl6^-/-^* mice show increased keratinocyte death after mechanical damage

Since histologic analysis of the skin after tape stripping revealed greater numbers of degenerated keratinocytes in *Elovl6^-/-^* mice than in wild-type mice (Figure 3A), we speculated that the number of dead cells were greater in the skin of *Elovl6^-/-^* mice than in that of wild-type mice after tape stripping. Indeed, flow cytometry analysis demonstrated that, although the proportion of dead keratinocytes in the epidermis of *Elovl6^-/-^* mice was comparable with that in wild-type mice in the steady state, tape stripping enhanced keratinocyte death in *Elovl6^-/-^* mice significantly more than in wild-type mice (Fig. 3B, C). These results suggest that Elovl6 suppressed mechanical damage-induced keratinocyte death.

**Fig. 3.**
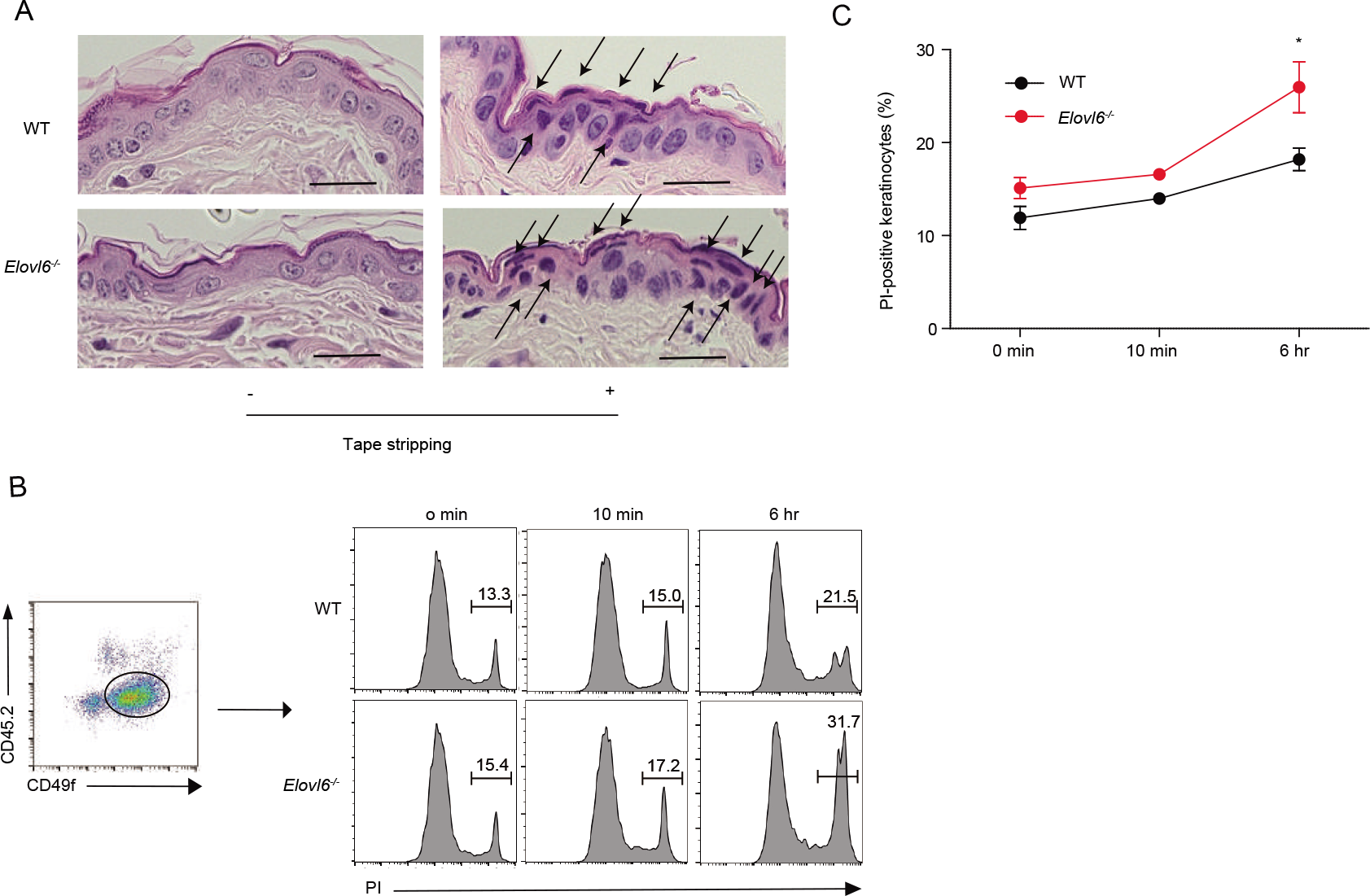
Elovl6 suppressed mechanical damage–induced keratinocyte death. (A) Representative histopathology of wild-type and *Elovl6^-/-^* mice 4 h after tape stripping. Black arrows indicate degenerated keartinocytes with irregularly shaped nuclei. Scale bar, 50 pm. (B, C) Flow cytometry of epidermal cells isolated from the skin of wild-type and *Elovl6^-/-^* mice at the indicated time points after tape stripping. Cells were stained with anti-CD45.2, anti-CD49f and propidium iodide (PI) and the proportion of PI+ cells in CD45.2-CD49f+ cells were shown (n = 6 to 10 per group). Error bars indicate 1 SD; *, *P*< 0.05. Data are representative of two independent experiments.

### Cis-vaccenic acid (CVA) is increased in *Elovl6^-/-^* mice

To investigate how keratinocytes death and the skin inflammation after tape stripping were exacerbated in *Elovl6^-/-^* mice, we analyzed the fatty acid composition of the epidermis of wild-type and *Elovl6^-/-^* mice. Unlike in our previous reports of increased PA levels in the lung and liver of *Elovl6^-/-^* mice (Matsuzaka *et al*. 2007; Sunaga *et al.* 2013), PA was not increased in the epidermis (Figure 4A). Instead, *Elovl6^-/-^* mice had significantly increased CVA levels (Figure 4A) and greater epidermal expression of the long-chain fatty acid elongases *Elovl1, Elovl3,* and *Elovl5* and of the stearoyl-CoA desaturase *Scd3* than did wild-type mice (Figure 4B). Among these, *Elovl5* and *Scd3* may influence CVA generation through the elongation of POA (C16:1n-7) (Burns *et al.* 2012) and by the conversion of PA to POA (Guillou *et al*. 2010), respectively (Figure supplement 4). These results suggest that CVA might be involved in the keratinocytes death and skin inflammation after tape stripping in *Elovl6^-/-^* mice.

**Fig. 4.**
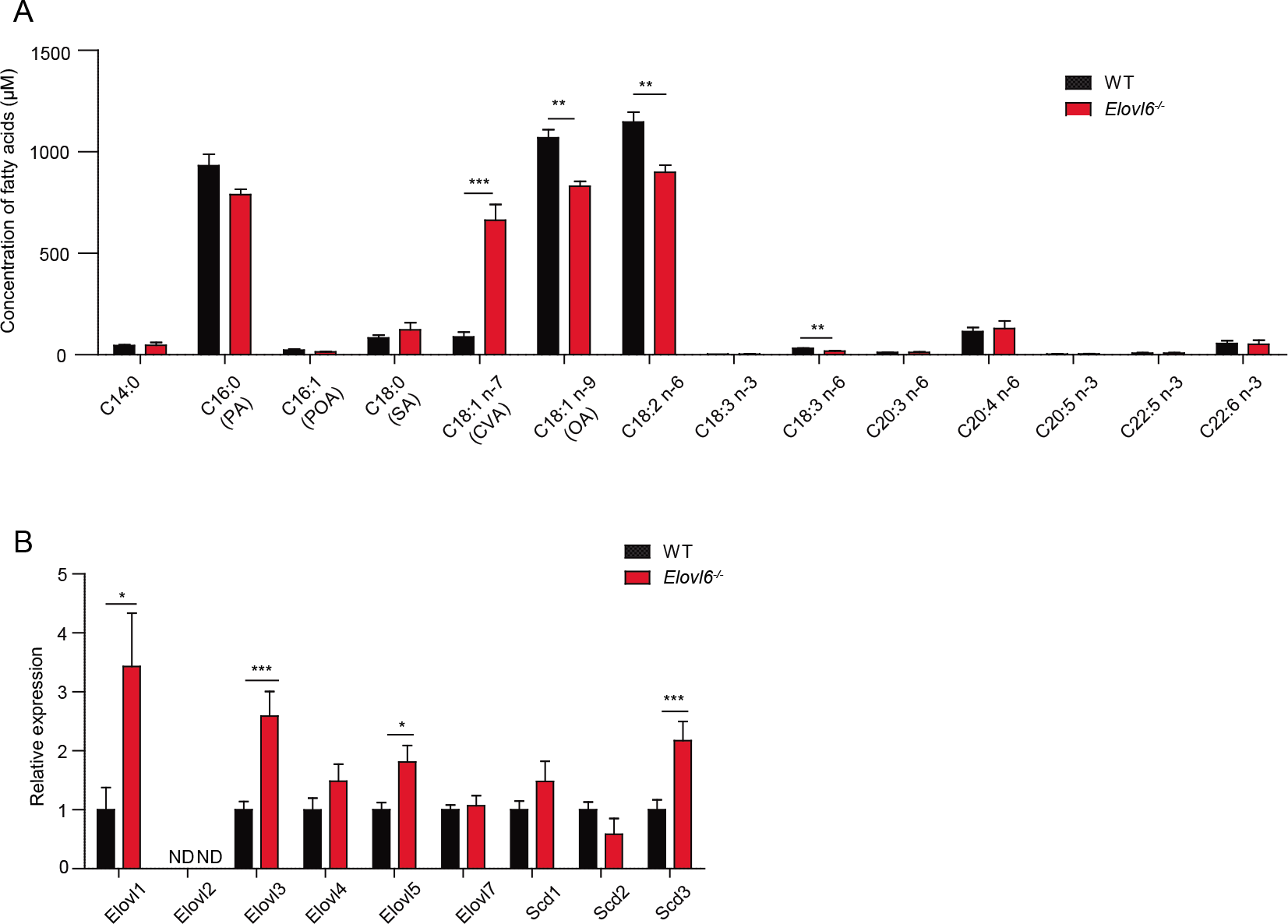
Elovló suppressed cis-vaccenic acid production in keratinocytes. (A) Fatty acid composition of epidermis in wild-type and *Elovl6^-/-^* mice (n = 4 per group). (B) Quantitative RT-PCR analysis of the epidermis for the expression of long-chain fatty acid elongases and stearoyl-CoA desaturases in wild-type and *Elovl6^-/-^* mice (n=10). Error bars indicate SD. ND, not done. *, *P*< 0.05; **, *P*< 0.01; \*\*\**P*<0.001.

### CVA induces keratinocyte death

To address whether CVA is involved in keratinocyte death, we cultured a human keratinocyte cell line HaCaT and primary keratinocytes derived from mice in the presence of CVA. We found that CVA decreased the numbers of live cells of HaCaT cells and primary keratinocytes in a dose-dependent manner (Figure 5A, B) and increased the proportion of dead primary keratinocytes (Figure 5C). In contrast, neither oleic acid (OA), PA, POA, SA, nor trans-vaccenic acid (TVA) influenced the number of live primary keratinocytes after culture (Figure 5A, B, D). In addition, CVA decreased the number of live peritoneal macrophages as well (Figure supplement 5A). CVA did not affect the proliferation of HaCaT cells but instead increased the number of dead cells compared with those after the addition of OA (Figure 5E), thus indicating that treatment with CVA induced cell death of HaCaT cells. This cell death was not affected by triacsin C, an inhibitor of long-chain acyl-CoA synthetases (Igal *et al*. 1997; Wang *et al*. 2012) (Figure supplement 5B), suggesting that CVA itself, but not its metabolites, induced death of HaCaT cells. Morphologic analyses under transmission electronic microscopy demonstrated increased plasma membrane rupture without in the keratinocytes after CVA treatment (Figure supplement 5C). In vivo, we found that topical application of CVA, but not OA, at a dose of 45 mM to the dorsal skin of wild-type mice increased the proportion of dead keratinocytes, as analyzed by flow cytometry (Figure 5F). Anti-cleaved caspase-9 (CC9) antibody did not stain CVA-treated dead keratinocytes (Figure supplement 5D). Together, these results suggest that CVA induced non-apoptotic cell death. Pretreatment with necrostatin-1 or necrosulfonamide, which are inhibitors of receptor-interacting protein 1 (RIP1) kinase and mixed lineage kinase domain-like protein (MLKL), respectively, did not suppress the CVA-induced death of keratinocytes (Figure supplement 5E), suggesting that the cell death due to CVA likely was not necroptosis (Skrzeczynska-Moncznik *et al*. 2015; Zhao *et al*. 2017). These combined results suggest that CVA induced necrosis rather than programmed cell death of keratinocytes. In addition, treatment with inhibitors of oxidative stress (IM-54) or cyclophilin D (cyclosporine A) did not influence the cell death (Figure supplement 5E), suggesting that the CVA-induced necrosis of keratinocytes was independent of oxidative stress (IM-54) or cyclophilin D–mediated changes in mitochondrial permeability (Chen *et al*. 2013; Nakagawa *et al*. 2005; Zeng *et al*. 2016).

**Fig. 5.**
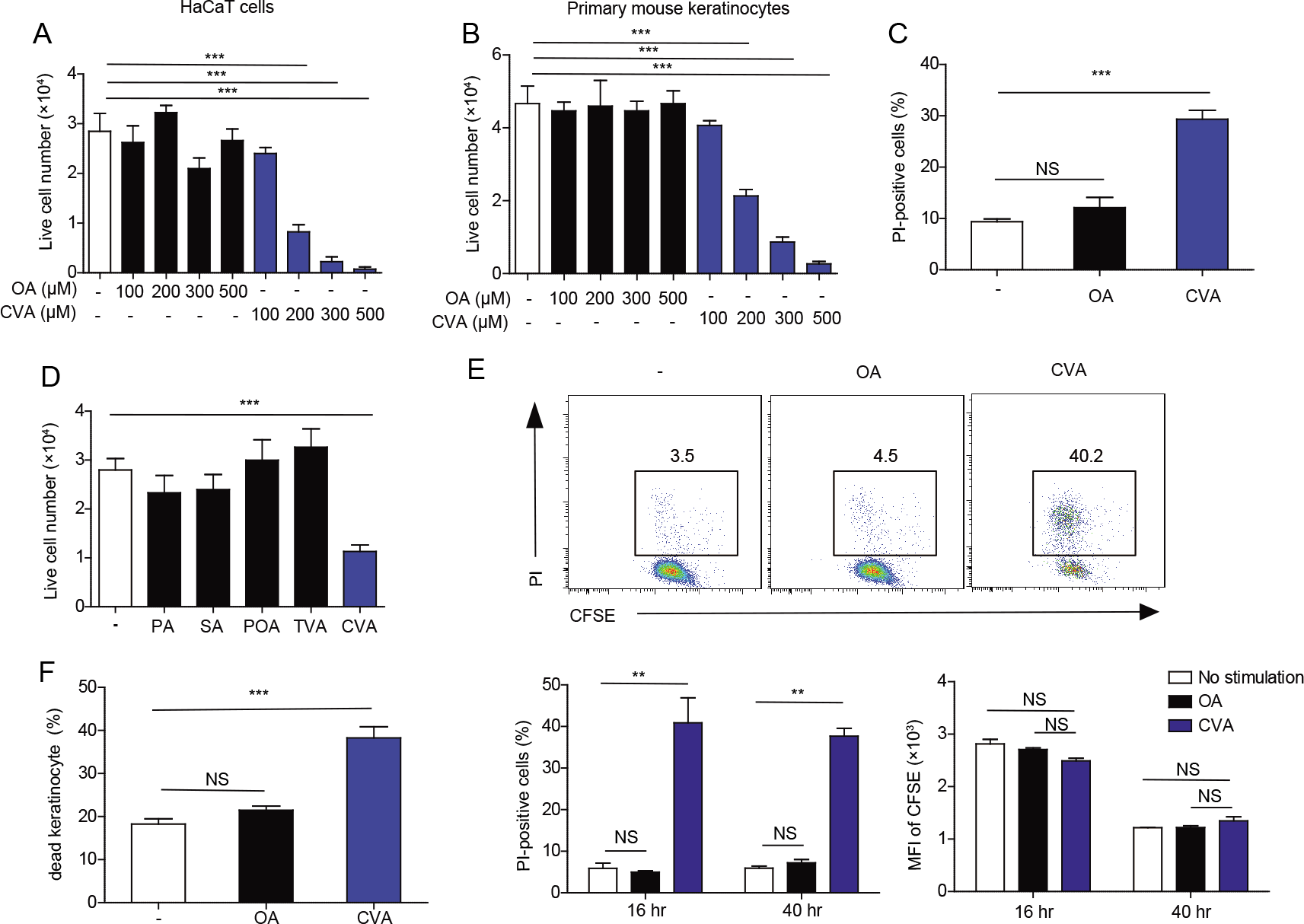
Cis-vaccenic acid (CVA) induces keratinocyte death in the skin. (A, B, D) Number of live cells in cultures of HaCaT cells (A) and primary keratinocytes (B, D) 16 h after stimulation with the indicated concentration of fatty acids (A, B) or with 300 μM of the indicated fatty acid (D) (n = 3 per group). (C) Primary keratinocytes were stimulated for 6 h with 300 μM of the indicated fatty acid and analyzed for propidium iodide (PI)-positive (i.e. dead) cells by flow cytometry (n = 4 or 5 per group). (E) HaCaT cells were labeled with 5-(and 6)-carboxyfluorescein diacetate succinimidyl ester (CFSE) and stimulated or not with 300 μM of OA or CVA for 16 h or 40 h. Cells were then stained with propium iodide (PI) and analyzed by flow cytometry. (**F**) Flow cytometry of epidermal cells isolated from the skin of wild-type mice that received topical application of ethanol (control) or 45mM of OA or CVA to the dorsal skin for 6 h (n = 6 per group). Error bars indicate SD. **, *P*< 0.01; ***, *P*< 0.001.

### CVA increased IL-1β and CXCL-1 production

Since CVA induced non-apoptotic cell death, we then examined whether CVA increased the release of DAMPs from dead cells. The addition of CVA, but not OA, to cultures of primary keratinocytes from wild-type mice increased the concentrations of HMGB-1 and IL-1α in the supernatants (Figure 6A). We further examined whether these DAMPs is involved in the increase in IL-1β and CXCL-1 expression. Stimulation of primary keratinocytes derived from wild type or *Elovl6^-/-^* mice in vitro and of the epidermis from the either genotype of mice in vivo with HMGB-1 or IL-1α induced *Il1β* and *Cxcl1*, and the expression levels of these cytokines transcripts did not differ between both genotypes of mice (Figure supplementary 6A, B). These results suggest that CVA enhanced IL-1β and CXCL-1 production by keratinocytes via HMGB-1 or IL-1α. Indeed, we found that topical application of CVA, but not OA, at a dose of 45 mM to the dorsal skin of wild-type mice increased the expressions of *Il1β* and *Cxcl1* in the epidermis (Figure 6B). Finally, treatment with either antagonist of IL-1 receptor or CXCR-2 intradermally and intraperitoneally reduced epidermal thickness and the number of neutrophils in the skin of *Elovl6^-/-^* mice (Figure 6C, D). These results suggest that the IL-1β and CXCL-1 produced by keratinocytes played crucial roles in the exacerbation of mechanical damage-induced skin inflammation in *Elovl6^-/-^* mice. Taken all together, these results suggest that tape stripping triggered keratinocyte death and release of HMGB-1 and IL-1α, which then stimulated the surrounding live keratinocytes to produce IL-1β and CXCL-1. Elovl6 deficiency increased the proportion of CVA in the skin, which accelerated keratinocyte death triggered by tape stripping and the subsequent signaling cascade to the production of IL-1β and CXCL-1, thus exacerbating dermatitis (Figure 6E).

**Fig. 6.**
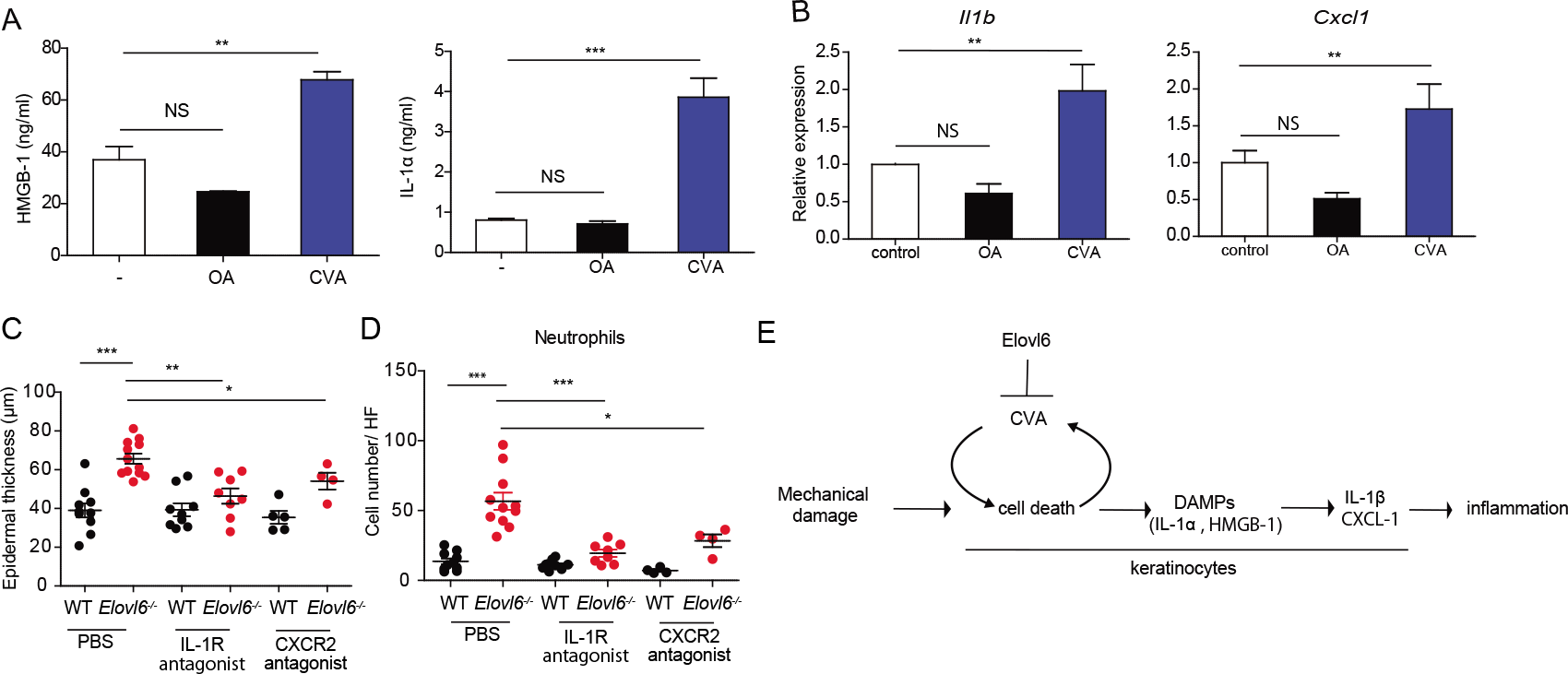
CVA increased IL-1β and CXCL-1 production. (A) Enzyme-linked immunosorbent assay of HMGB-1 (n = 4 per group) and cytokine bead array of IL-1α (n = 3 per group) in the supernatant of cultured primary keratinocytes 10 h after initiation of stimulation with 300 μM OA or CVA. (B) Quantitative RT-PCR analysis of *Il1β* and *Cxcl1* in the epidermis of wild-type mice 6 h after topical application of ethanol (control) (n = 10) or 15mM of OA (n = 13) or CVA (n = 14) (B). (C, D) Wild-type and *Elovl6^-/-^* mice were treated with PBS (n = 10 and 12, respectively), an IL-1 receptor antagonist (n = 9 and 8, respectively), or a CXCR-2 antagonist (n = 5 and 4, respectively) daily for 9 days, from the beginning on the day of tape stripping. Epidermal thickness (C) and the number of infiltrating neutrophils (D) were analyzed on day 9. (E) A proposed signal pathway from mechanical damage onto the skin to skin inflammation. Error bars indicate SD; *, *P* < 0.05; **, *P* < 0.01, ***, *P* < 0.001; NS, not significant. Data are representative of at least two independent experiments.

## Discussion

Previous studies have revealed increased PA and decreased OA contents in the liver and lung of *Elovl6^-/-^* mice compared with wild-type mice (Matsuzaka *et al*. 2007; Sunaga *et al*. 2013). In the current study, we noted that the OA level in the skin of *Elovl6^-/-^* mice was decreased compared with that in wild-type mice. However, PA content did not differ between the two genotypes. Instead, epidermal levels of CVA were greater in *Elovl6^-/-^* mice than in wild-type mice, presumably owing to its efficient conversion to CVA by the concomitant induction of SCD3 and Elovl5 productions (Burns *et al*. 2012; Guillou *et al*. 2010). Although it remains unclear at present how Elovle6 regulates the fatty acid composition and the enzyme alteration in the skin, these combined results suggest that regulation of the elongation of saturated and monounsaturated fatty acids is—in part—dependent on the organs or tissues.

The biologic function of CVA has been poorly understood. Here, we demonstrated that CVA directly induced cell death in cultures of primary keratinocytes from mice. Specifically, the cells killed by CVA lacked one of the hallmarks of apoptosis, namely caspase-9 activation. In addition, morphologic analyses revealed characteristics of necrosis, including plasma membrane rupture without blebbing (Krysko *et al*. 2008). None of the inhibitors of necroptosis, oxidative stress, or cyclophilin-D-associated cell death—including necrostatin-1, NSA, IM-54, and cyclosporine A—inhibited the CVA-induced cell death. In addition, triacsin-C treatment failed to suppress CVA-induced cell death. These results suggest that CVA induced necrosis rather than programmed cell death of keratinocytes.

Long-chain fatty acids elicit a variety of biologic effects, including cell death. For example, PA induces apoptosis in many cell types, and this response is abrogated by OA (Gillet *et al*. 2015; Sunaga *et al*. 2013). In contrast, PA induces RIP1/RIP3-dependent necroptosis in RAW 264.7 cells (Kim *et al*. 2017), and (although the results are controversial) OA has been reported to induce the death of various cell types (Brinkmann *et al*. 2013; Moravcova *et al*. 2015). The molecular mechanisms of these cytotoxic effects remain poorly understood; features proposed to be involved in these toxicities include loss of membrane integrity, changes in mitochondrial transmembrane potential, activation of caspase-3, and the production of reactive oxygen species (Brinkmann *et al*. 2013; Fontana *et al*. 2013). In the current study, we found that, beginning soon after its addition to the cultures, CVA was severely cytotoxic to HaCaT cells, mouse peritoneal macrophages, and primary keratinocytes; CVA can thus be added to the list of possible lipotoxins.

Previous studies have demonstrated that, compared with OA and SA, supplementation with both CVA and TVA (dose, 30 μM) significantly suppress the growth of HT-29 tumor cells after culture for 9 days (Awad *et al*. 1995). Moreover, CVA leads to greater hydrolysis of phosphoinositides in the plasma membrane than does TVA (Awad *et al*. 1995), suggesting that CVA is incorporated into the plasma membrane and affects the phospholipids composition. In the present study, we showed that CVA at concentrations of 200 µM or greater induced the death, rather than the suppression of growth, of keratinocytes, thus suggesting that the higher amount of CVA induces damage of the plasma membrane sufficiently to induce necrosis. In addition, given that TVA did not induce keratinocyte death, the cytotoxic effect of CVA may be structure dependent. Whereas trans-unsaturated fatty acids have a linear structure and can be packed regularly in the plasma membrane, cis-unsaturated fatty acids such as OA and CVA, which have a characteristic angular kink, may distort the structure of the lipid bilayer and thus destabilize the plasma membrane (Fontana *et al*. 2013). Therefore, although further studies are required to determine the detailed mechanism of CVA-induced cell death, we speculate that incorporation of CVA into the plasma membrane creates a bulky 3-dimensional structure compared with those associated with other cis-monounsaturated fatty acids and thus induces cell death by disrupting the plasma membrane.

Fatty acids reportedly play important roles in modulating dermatitis. For example, a high-fat diet enriched with oleic acid impairs contact hypersensitivity responses to trinitrochlorobenzene and FITC (Katagiri *et al*. 2008). In addition, oral administration of docosahexaenoic acid leads to the generation of regulatory T cells, which thus attenuate dinitrochlorobenzene-induced dermatitis (Han *et al*. 2015). Moreover, topical or oral application of linoleic acid and TVA, which are enriched in milk fat, decreases the severity of OVA-induced atopic dermatitis (Sun *et al*. 2011).

Atopic dermatitis (AD) is one of the most common skin diseases and is characterized by pruritic and eczematous skin lesions. The outermost layer of the epidermis, the stratum corneum, contains decreased levels of ceramides (very long-chain fatty acids), leading to impaired barrier function in patients with AD, and the average chain length of ceramide fatty acids is negatively correlated with epidermal permeability in these patients (Ishikawa *et al*. 2010). Moreover, *Elovl1^-/-^* and *Elovl4^-/-^* mice, both of which demonstrate a global decrease in the chain length of ceramide fatty acids of the stratum corneum, show impaired barrier function (Li *et al*. 2007; Sassa *et al.* 2013). Mechanical stress, such as scratching, increases the severity of AD by removing the stratum corneum (thus diminishing the epidermal barrier function) and by inducing the production of pro-inflammatory cytokines (Verhoeven *et al*. 2008; Wahlgren 1999). On the other hand, psoriasis is characterized by well-demarcated scaly erythema and plaque, which sometimes show itching, and histopathologically, these lesions reveal hyperproliferation of keratinocytes and neutrophil infiltration (Hirotsu *et al*. 2012). The skin lesions of psoriasis are well-known to be triggered or exacerbated, as the Koebner phenomenon, by mechanical stress (Köbner 1876). Our current results suggest that Elovl6 may regulates mechanical stress-induced exacerbation of skin inflammation due to inhibition of keratinocyte death by CVA in patients with AD and psoriasis.

## Materials and Methods

### Mice

*Elovl6^-/-^* mice on the C57BL/6J background were described previously (Matsuzaka *et al*. 2007). C57BL/6J mice raised under specific pathogen-free conditions were purchased from Clea Japan (Tokyo, Japan). Germ-free mice were bred and maintained in isolators at Sankyo Laboratories (Tsukuba, Japan), and their germ-free status was routinely confirmed by in-house aerobic and anaerobic culture of feces. *K14*-Cre, *Ticam1^-/-^*, and *Myd88^-/-^* mice on the B57BL/6 background were purchased from Jackson Laboratories (Bar Harbor, Maine, USA). *Elovl6^fl/fl^* mice were crossed with K14-Cre transgenic mice to generate Elovl6-knockout mice specifically in keratinocytes (*Elovl6^fl/fl^ K14-Cre*-Cre). Mice between 8 and 10 weeks of age were used for the experiments. All experiments were performed in accordance with the guidelines of the animal ethics committee of the University of Tsukuba Animal Research Center.

### Tape stripping (Nakajima *et al.* 2012)

To generate mechanical damage-induced dermatitis, a 2.5×2.5 cm area of the dorsal skin was shaved and tape-stripped 20 times by using adhesive tape (Johnson and Johnson); a 1×1-cm piece of sterile gauze moistened with 100 μl PBS was placed on the shaved skin and secured with transparent bio-occlusive tape (Tegaderm Roll, 3M, Maplewood, Minnesota, USA) to prevent the mice from licking the area. These procedures were repeated every other day until analysis.

### Cytokine measurement of epidermis or cultured keratinocytes

Dorsal skin samples before and after tape stripping were resected from adult mice and incubated in RPMI medium in the presence of dispase II (3 mg/ml) (Wako Pure Chemical, Osaka, Japan) for 1 h at 37 °C under 5% CO_2_. The epidermis was then separated from the dermis under a stereomicroscope. Samples of epidermis (diameter, 4 mm) were cultured in 50 μl of DMEM containing 10% FBS in a 96-well plate at 37 °C under 5% CO_2_ for 24 h and the concentrations of IL-1β, CXCL-1, TNFα, and IL-10 in the culture supernatants were measured by using cytometric bead arrays (CBA) (BD Biosciences) according to the manufacturer’s protocol. The skin of new born mice (younger than 3 days old) was incubated in CnT-07 medium (CELLnTEC Advanced Cell Systems) in the presence of dispase II (1mg/ml) (CELLnTEC Advanced Cell Systems) at 4 °C for 16 h. The epidermis was isolated from the skin and incubated in Accutase (CellnTEC Advanced Cell Systems) at room temperature for 20 min. The keratinocytes collected were maintained in CnT-07 medium (CELLnTEC Advanced Cell Systems) according to the manufacturer’s protocol. Keratinocytes were then stimulated with long-chain fatty acids at 37 °C under 5% CO_2_ for 10 h and the culture supernatants were analyzed for IL-1α and HMBG-1 by flow cytometry using cytokine beads array (CBA) and for HMGB-1 using an ELISA KIT II (Shino-test Corporation). Keratinocytes were also stimulated with IL-1α or HMGB-1 at 37 °C under 5% CO_2_ for 3 h and analyzed for *Il1b* and *Cxcl1* by quantitative real-time PCR analysis (qRT-PCR).

### qRT-PCR

Total RNA was isolated by using ISOGEN (Wako Pure Chemical). Quantitative real-time PCR analysis was performed on a 7500 Fast Real-Time PCR System (Applied Biosystems, Foster City, California, USA) with Power SYBR Green PCR Master Mix (Applied Biosystems). Results are presented relative to those of the housekeeping gene encoding GAPDH (*gapdh*). Primers used were as follows: *Il1b* fwd, GAAATGCCACCTTTTGACAGTG; *Il1b* rev, TGGATGCTCTCATCAGGACAG; *Cxcl1* fwd, CTGGGATTCACCTCAAGAACATC; *Cxcl1* rev, CAGGGTCAAGGCAAGCCTC; *Ccl2* fwd, CAGGTCCCTGTCATGCTTC; *Ccl2* rev ATGAGTAGCAGCAGGTGAGTG; *Elovl6* fwd, ACAATGGACCTGTCAGCAAA; *Elovl6* rev, GTACCAGTGCAGGAAGATCAGT; *Tnfa* fwd, CCTGTAGCCCACGTCGTAG; *Tnfa* rev, GGGAGTAGACAAGGTACAACCC; *Il1a* fwd, AGGGAGTCAACTCATTGGCG; *Il1a* rev, TGGCAGAACTGTAGTCTTCGT; *Il6* rev, TCCACGATTTCCCAGAGAAC; *Il10* fwd, GCTGGACAACATACTGCTAACC; *Il10* rev, ATTTCCGATAAGGCTTGGCAA; *Il33* fwd, GGTGAACATGAGTCCCATCA; *Il33* rev, CGTCACCCCTTTGAAGCTC; *Tgfb* fwd, TGACGTCACTGGAGTTGTACGG; *Tgfb* rev, GGTTCATGTCATGGATGGTGC; *Ccl2* rev ATGAGTAGCAGCAGGTGAGTG; *Ifng* fwd, ACAGCAAGGCGAAAAAGGATG; *Ifng* rev, TGGTGGACCACTCGGATGA; *Il4* fwd; ATCATCGGCATTTTGAACGAGG; *Il4* rev; TGCAGCTCCATGAGAACACTA; *Il17* fwd, TTTAACTCCCTTGGCGCAAAA; *Il17* rev, CTTTCCCTCCGCATTGACAC; *Elovl1* fwd, TCCAAAGCTACCCTCTGATGG; *Elovl1* rev, AGGGAGAGTATCACCAGTGAGA; *Elovl2* fwd, ACGCTGGTCATCCTGTTCTT; *Elovl2* rev, GCCACAATTAAGTGGGCTTT; *Elovl3* fwd, TTCTCACGCGGGTTAAAAATGG; *Elovl3* rev, GAGCAACAGATAGACGACCAC; *Elovl4* fwd, GCCCTGTGGTGGTATTTTGT; *Elovl4* rev, TGGTGGTACACGTGAAGGAA; *Elovl5* fwd, GGTGGCTGTTCTTCCAGATT, *Elovl5* rev, CCCTTCAGGTGGTCTTTCC, *Elovl6* fwd, ACAATGGACCTGTCAGCAAA; *Elovl6* rev, GTACCAGTGCAGGAAGATCAGT; *Elovl7* fwd, CATCGAGGACTGTGCGTTTTT; *Elovl7* rev, CCAGGATGATGGTTTGTGGCA; *Scd1* fwd, TCAACTTCACCACGTTCTTCA; *Scd1* rev, CTCCCGTCTCCAGTTCTCTT; *Scd2* fwd, TGGTTTCCATGGGAGCTG; *Scd2* rev, TTGATGTGCCAGCGGTACT; *Scd3* fwd, CTGACCTGAAAGCCGAGAAG; *Scd3* rev, GCAGAATGCCAGGCTTGTA; *Gapdh* fwd, AGGTCGGTGTGAACGGATTTG; and *Gapdh* rev TGTAGACCATGTAGTTGAGGTCA.

### Histology

For histologic analysis, mouse skin was fixed in 10% formalin, embedded in paraffin, sectioned, and stained with hematoxylin and eosin. For analysis of epidermal thickness or cell number, 18 randomly selected sites were evaluated by using light microscopy or fluorescent microscopy (MZ-X710, Keyence, Osaka, Japan) and its associated software.

### Analysis of cleaved caspase-9

Cells were treated or not with CVA or UV (180 mJ/cm^2^ for 3 min) and then fixed with 4% paraformaldehyde for 15 min and then with 5 % BSA in PBS containing 0.3% Triton X-100 for 1 h. Subsequently, cells were incubated with rabbit anti-cleaved caspase-9 antibody (1:200; Cell Signaling) for 1 h at room temperature, followed by incubation with Alexa Fluor 594-conjugated donkey anti-rabbit IgG (1:200; Thermo Fisher Scientific) secondary antibody for 30 min. Finally, cells were counterstained with DAPI.

### Cell death and proliferation analyses

Primary keratinocytes from neonatal epidermis, prepared as described above, or human HaCaT cells were maintained in CnT-07 medium, as described above, and DMEM with 10% FBS, respectively, at 37 °C under 5% CO_2_. Peritoneal macrophages were harvested by lavage of the peritoneal cavity and suspended in DMEM containing 10% FBS. Primary keratinocytes and HaCaT cells and peritoneal macrophages were seeded onto 48-well plates at a density of 1×10^5^ cells and 2×10^5^ cells/ well, respectively. After incubation at 37 °C for 2 h, the cells were washed with PBS three times to remove unattached cells and stimulated with different concentrations of free fatty acids dissolved in ethanol (final dose of 100 ∼ 500 μM), including oleic acid (OA), palmitic acid (PA), stearic acid (SA), palmitoleic acid (POA) (Wako Pure Chemical), trans-vaccenic acid (TVA) or cis-vaccenic acid (CVA) (Sigma-Aldrich). To block specific types of cell death or inhibit long-chain acyl-coenzyme A synthetase, cells were pretreated with 1 mM of necrostatin-1 (Cayman Chemicals), 1 mM of necrosulfonamide (Cellagen Technology), 2 mM of IM-54 (Sigma Aldrich), 1 mM of cyclosporine A (Sigma Aldrich), or 10 μM of triacsin C (Abcam) for 6 h before fatty acid stimulation.

For proliferation assay, these cells were stained or not with 10 μM of CFSE (Invitrogen) for 5min at 37°C before fatty acid stimulation, according to the manufacturer instructions. Cells were then stained with propidium iodine (PI) and analyzed for PI-positive and -negative cell populations and CFSE dilution by flow cytometry. For the analysis of cell death in vivo, skin tissue was incubated in 0.5% trypsin (Wako) in PBS, and separated into epidermis. Epidermal cells were stained with CD45.2 (clone:104, BD pharmingen), CD49f (clone: GoH3, Miltenyi Biotec) and PI, and then analyzed by flow cytometry.

### Transmission electron microscopy

Cultured keratinocytes were fixed by incubating 2.5% glutaraldehyde in PBS (pH 7.4) at 4 °C overnight, postfixed in 1% osmium tetraoxide at 4 °C for 30 min, and then dehydrated through graded concentrations of ethanol. Cells were then transferred to propylene oxide and embedded (Poly/Bed 812, Polysciences, Warrington, Pennsylvania, USA). The samples were analyzed by electron microscopy (JEM-1400, JEOL, Peabody, Massachusetts, USA).

### Cytokine stimulation

Primary mouse keratinocytes were stimulated with bovine HMGB-1 (Chondrex, Redmond, Washington, USA) or mouse IL-1α (Miltenyi Biotec, Bergisch Gladbach, Germany). For stimulation of keratinocytes in vitro, primary keratinocytes were stimulated 500ng/ml of bovine HMGB-1 or mouse IL-1α. For stimulation of keratinocytes in vivo, 200 ng of bovine HMGB-1 or mouse IL-1α in 50 µl PBS was injected intradermally.

### Fatty acid composition

Lipids from mouse epidermis were extracted by using the method of Bligh and Dyer (Breil *et al*. 2017). In brief, epidermis was extracted with chloroform/ methanol (1:2, v/v) solution. One molar NaCl solution and chloroform were added to break the monophase and incubated on ice for 10 min. After centrifugation at 300 G for 5 min, aqueous solution was discarded and the phase of chloroform was evaporated using nitrogen gas. Following the addition of acetonitrile/ 6N HCl (90/10, v/v), samples were incubated at 100 °C for 45 min. Finally, liquid-liquid extraction (Milne and Zhitomirsky 2018) with ethyl acetate was performed and the reconstituted samples were injected into an optimized LC/ MS/MS system. The relative abundance of each fatty acid was quantified by gas chromatography.

### Transepidermal water loss (TEWL) test

Fetuses obtained at E13.5 and E17.5 were incubated in methanol for 5 min and rinsed in PBS, followed by incubation with 0.1% toluidine blue for 24 h. After staining, fetuses were rinsed with PBS and the staining intensity was evaluated (Sassa *et al*. 2013). TEWL was also measured on the dorsal skin of wild-type and *Elovl6^-/-^* mice between 8 and 10 weeks of age by Tewameter^®^ TM 300 (Integral) before and after tape stripping (Sassa *et al*. 2013). Measurements were performed in triplicate for each mouse.

### Antagonist treatment

To neutralize the IL-1 receptor, mice received 200 µl of the IL-1 receptor antagonist anakinra (10 mg/ml) (Kineret, Swedish Orphan Biovitrum) intradermally and 300 µl of the same concentration of the antagonist intraperitoneally daily during the induction of dermatitis by the mechanical stress or OVA treatment. To block CXCR-2, mice received 200 µl of a CXCR-2 antagonist (0.25 mg/ml in PBS containing 1% DMSO) (SB225002, Cayman Chemical, Ann Arbor, Michigan, USA) intradermally and 150 µg of the same antagonist (0.5 mg/ml in PBS containing 1% DMSO) intraperitoneally daily during the induction of dermatitis by the mechanical stress.

Statistical analyses. Statistical analyses were performed by using an unpaired, two-tailed Student’s *t*-test (GraphPad Prism 6, GraphPad Software, La Jolla, USA). A *P* value less than 0.05 was considered to be statistically significant.

## Acknowledgments

We thank S. Mitsuishi and Y. Nomura for their secretarial assistance.

## Supplementary Figures

**Fig. S1.**
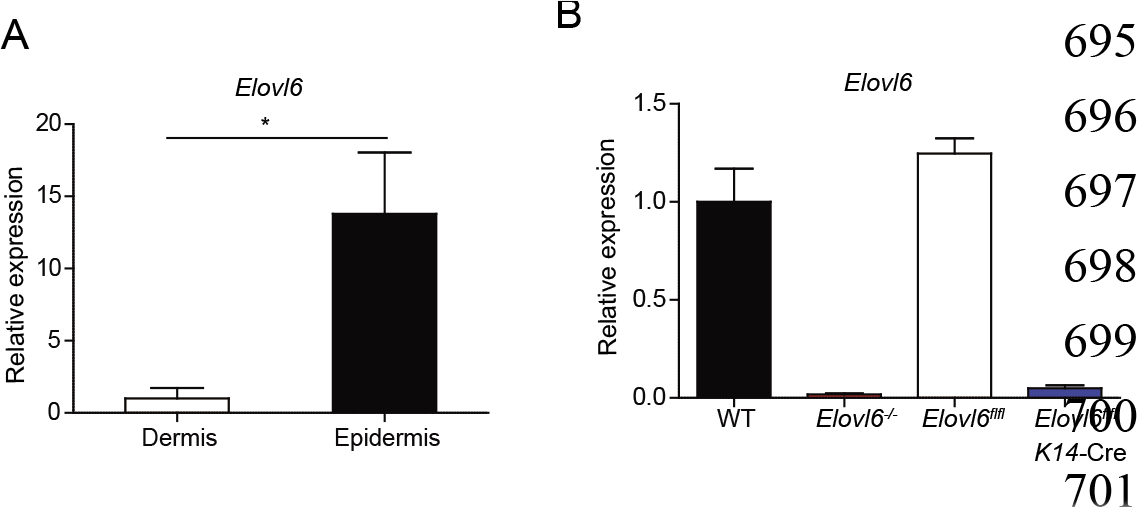
Expression of *Elovl6*. Quantitative RT-PCR analysis of *Elovl6* from dermis and epidermis (n = 3 per group) (**A**) and from the epidermis of wild-type (WT), *Elovl6^-/-^*, *Elovl6^flfl^*, *Elovl6*^fl/fl^ *K14*-Cre mice (n=3 to 5 in each group) (**B**). Error bars indicate SD. \**P*<0.05. Data are representative of more than two independent experiments.

**Fig. S2.**
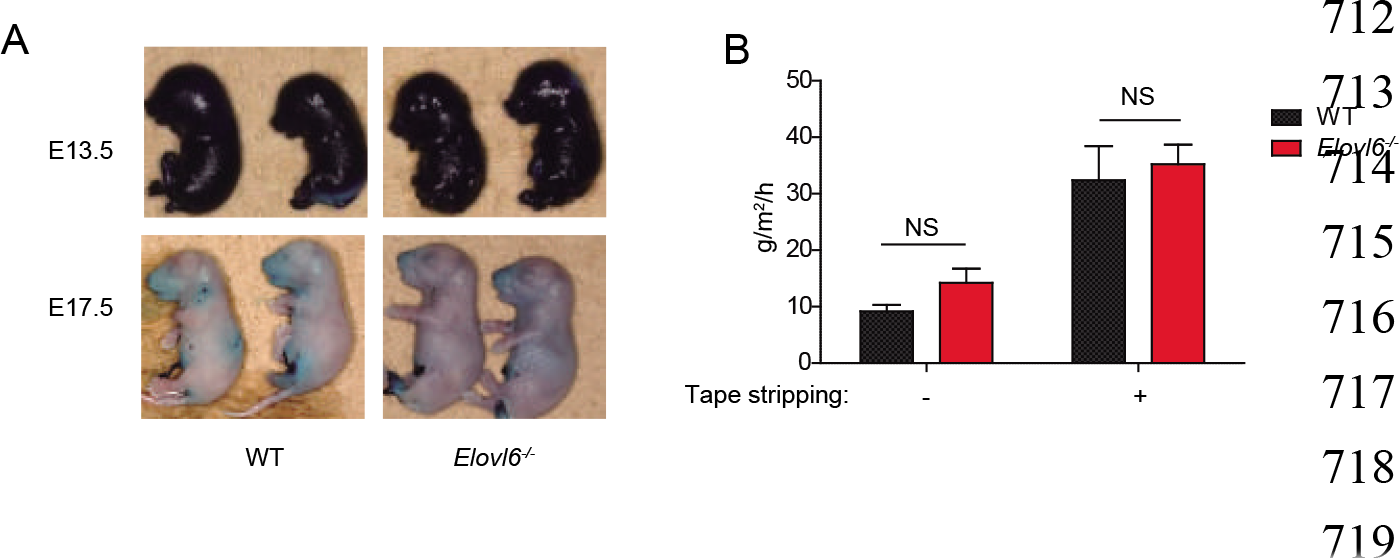
Comparable skin permeability barrier function between wild-type (WT) and *Elovl6^-/-^* mice. (**A**) The fetuses at E13.5 and E17.5 from WT and *Elovl6^-/-^* mice were stained with 0.1% toluidine blue for 24 h and photographed. (**B**) The transepidermal water loss of 6-8 weeks old WT and *Elovl6^-/-^* mice was measured before and after tape stripping (n = 14). Error bars indicate SD. NS, not significant. Data are representative of three independent experiments.

**Fig. S3.**
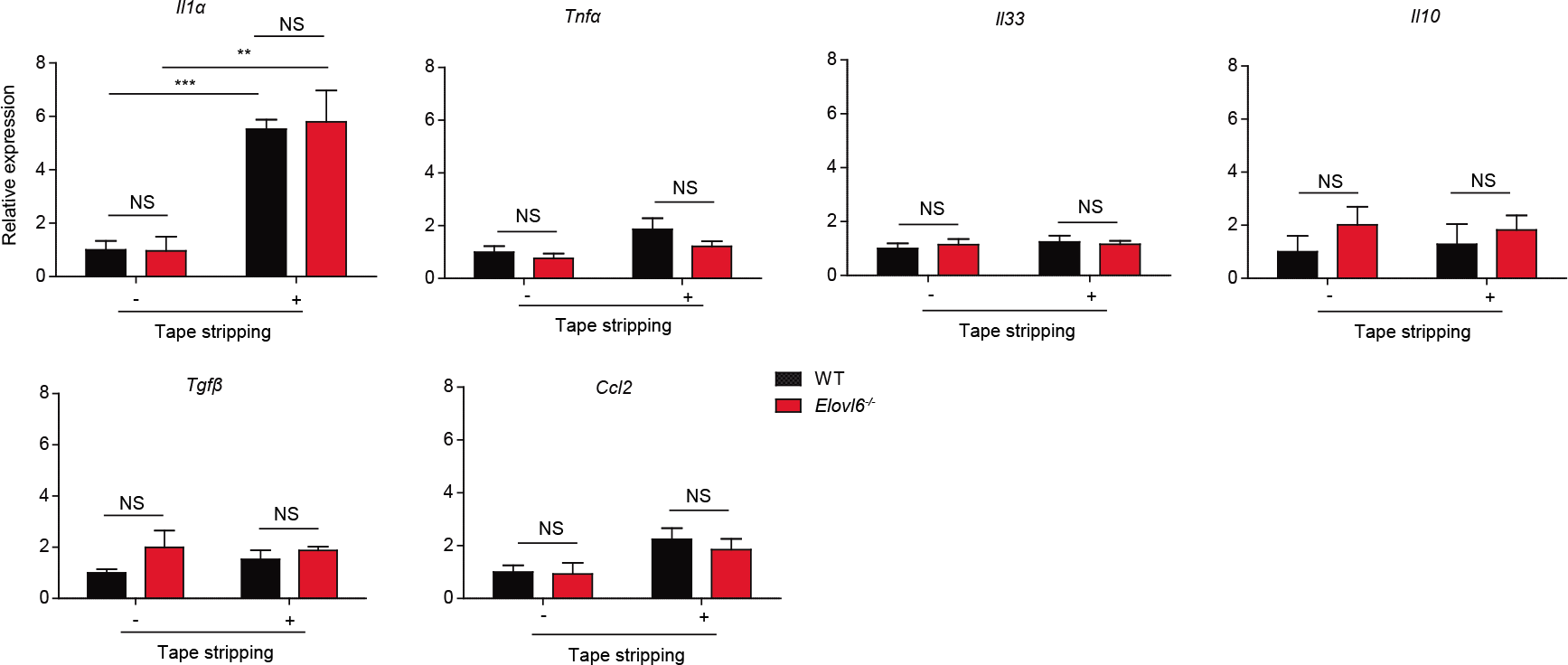
Expression of cytokines. Quantitative RT-PCR analysis of the epidermis isolated before and 12 h after tape stripping from WT and *Elovl6^-/-^* mice and cultured for 24 h (n = 11 in each group). Error bars indicate SD. NS, not significant. \*\**P*<0.01, \*\*\**P*<0.001. Data are representative of two independent experiments.

**Fig. S4.**
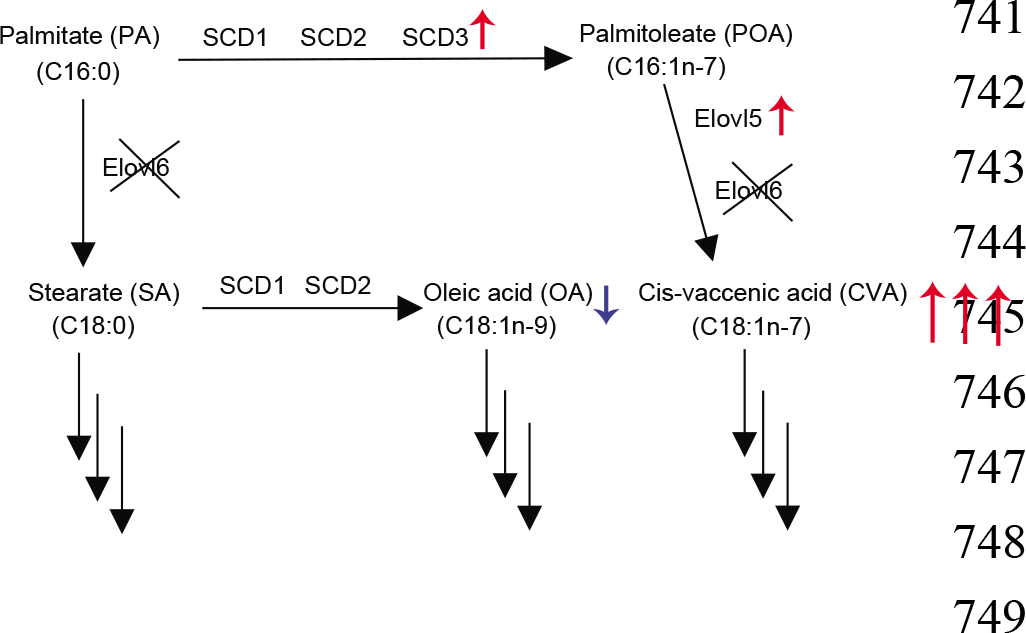
Schematic representation of the proposed pathway (black arrows) controlling cis-vaccenic acid (CVA) generation in *Elovl6^-/-^* keratinocyte. Red and blue arrows indicate the increase in SCD3, Elovl5, and CVA and the decrease in OA, respectively.

**Fig. S5.**
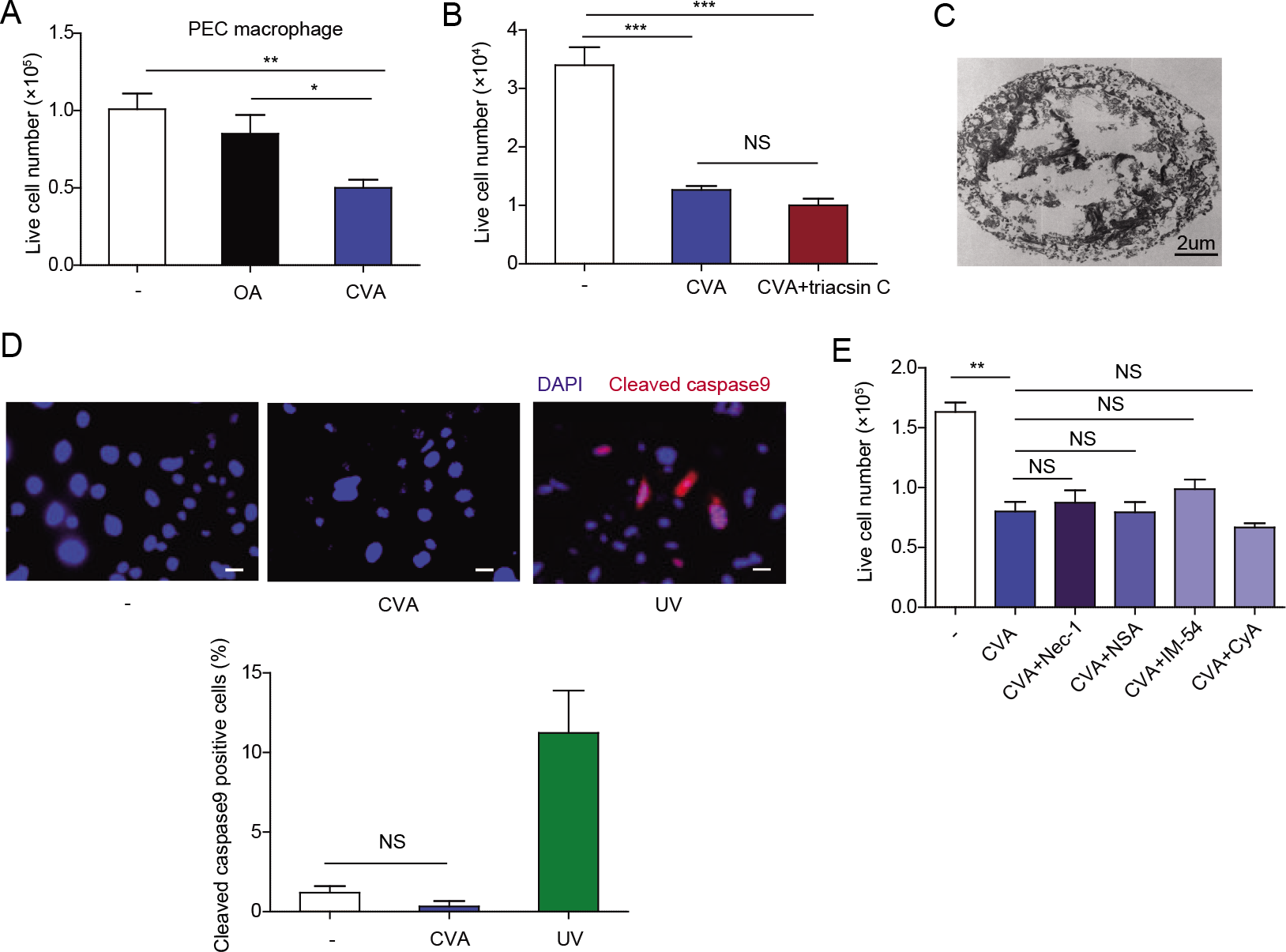
CVA induced non-programmed cell death. (A) Live cell number of primary peritoneal macrophages 16 h after stimulation with oleic acid (OA) or CVA (300 µM) (n = 3 in each group). (B, E) Primary mouse keratinocytes were cultured for 6 h in the presence or absence of 10 μM of triacsin C (B), or 1 mM necrostatin (Nec-1), 1 mM necrosulfonamide (NSA), 2 mM IM-54, or 1 mM cyclosporine (CyA) (E), followed by stimulation by adding 300 μM CVA; live cells were counted 16 h afterward (n = 3). (C) A representative dead primary keratinocyte induced by stimulation with 300 μM CVA for 10 h under a transmission electron microscope. (D) Immunofluorescence microscopic study of primary keratinocyte 10 h after stimulation with CVA or 6 h after ultraviolet irradiation. Cells were stained with anti-cleaved caspase 9, followed by Alexa Fluor 594-conjugated secondary antibody and DAPI. White bars indicate a scale (20 μm). Percentage of cleaved caspase 9-positive cells was calculated (n = 3). Error bars indicate SD. NS, not significant. *, *P*< 0.05; **, *P*< 0.01; ***, *P*< 0.001. Data are representative of more than two independent experiments.

**Fig. S6.**
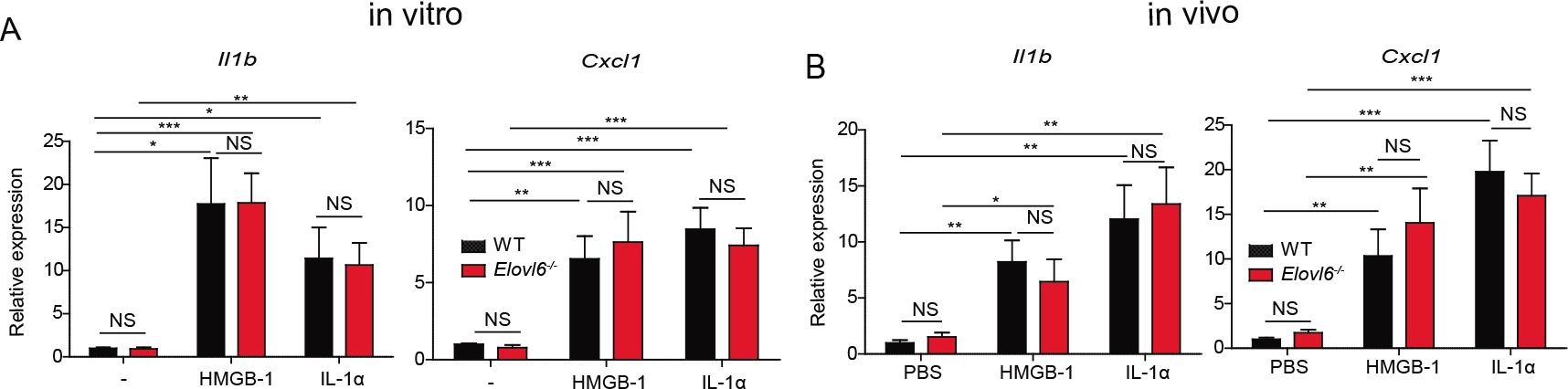
DAMPs increased IL-1β and CXCL-1 production. Quantitative RT-PCR analysis of *Il1β* and *Cxcl1* in keratinocytes of wild-type and *Elovl6^-/-^* mice after stimulation or not with HMGB-1 or IL-1α in vitro (n=10 per group) (A) and in the epidermis isolated 4 h after injection intradermally with PBS, HMGB-1, or IL-1α (n = 8 per each group) (B).

